# A small number of point mutations confer formate tolerance in *Shewanella oneidensis*

**DOI:** 10.1101/2024.06.28.601276

**Authors:** Megan C Gruenberg Cross, Elhussiny Aboulnaga, Michaela A TerAvest

## Abstract

Microbial electrosynthesis (MES) is a sustainable approach to chemical production from CO_2_ and clean electricity. However, limitations in electron transfer efficiency and gaps in understanding of electron transfer pathways in MES systems prevent full realization of this technology. *Shewanella oneidensis* could serve as an MES biocatalyst because it has a well-studied, efficient transmembrane electron transfer pathway. A key first step in MES in this organism could be CO_2_ reduction to formate. However, wild-type *S. oneidensis* does not tolerate high levels of formate. In this work, we created and characterized formate-tolerant strains of *S. oneidensis* for further engineering and future use in MES systems through adaptive laboratory evolution. Two different point mutations in a sodium-dependent bicarbonate transporter and a DUF2721-contianing protein separately confer formate tolerance to *S. oneidensis*. The mutations were further evaluated to understand their role in improving formate tolerance. We also show that the wild-type and mutant versions of the *S. oneidensis* sodium-dependent bicarbonate transporter improves formate tolerance of *Zymomonas mobilis*, indicating the potential for the transfer of this formate tolerance phenotype to other organisms.

**Importance:** *Shewanella oneidensis* is a bacterium with a well-studied, efficient extracellular electron transfer pathway. This capability could make this organism a suitable host for microbial electrosynthesis using CO_2_ or formate feedstocks. However, formate is toxic to *S. oneidensis*, limiting the potential for its use in these systems. In this work, we evolve several strains of *S. oneidensis* that have improved formate tolerance and we investigate some mutations that confer this phenotype. The phenotype is confirmed to be attributed to several single point mutations by transferring the wild-type and mutant versions of each gene to the unmutated strain. Finally, the formate tolerance mechanism of one mutation is studied using structural modeling and expression in another host. This study therefore presents a simple method for conferring formate tolerance to bacterial hosts.

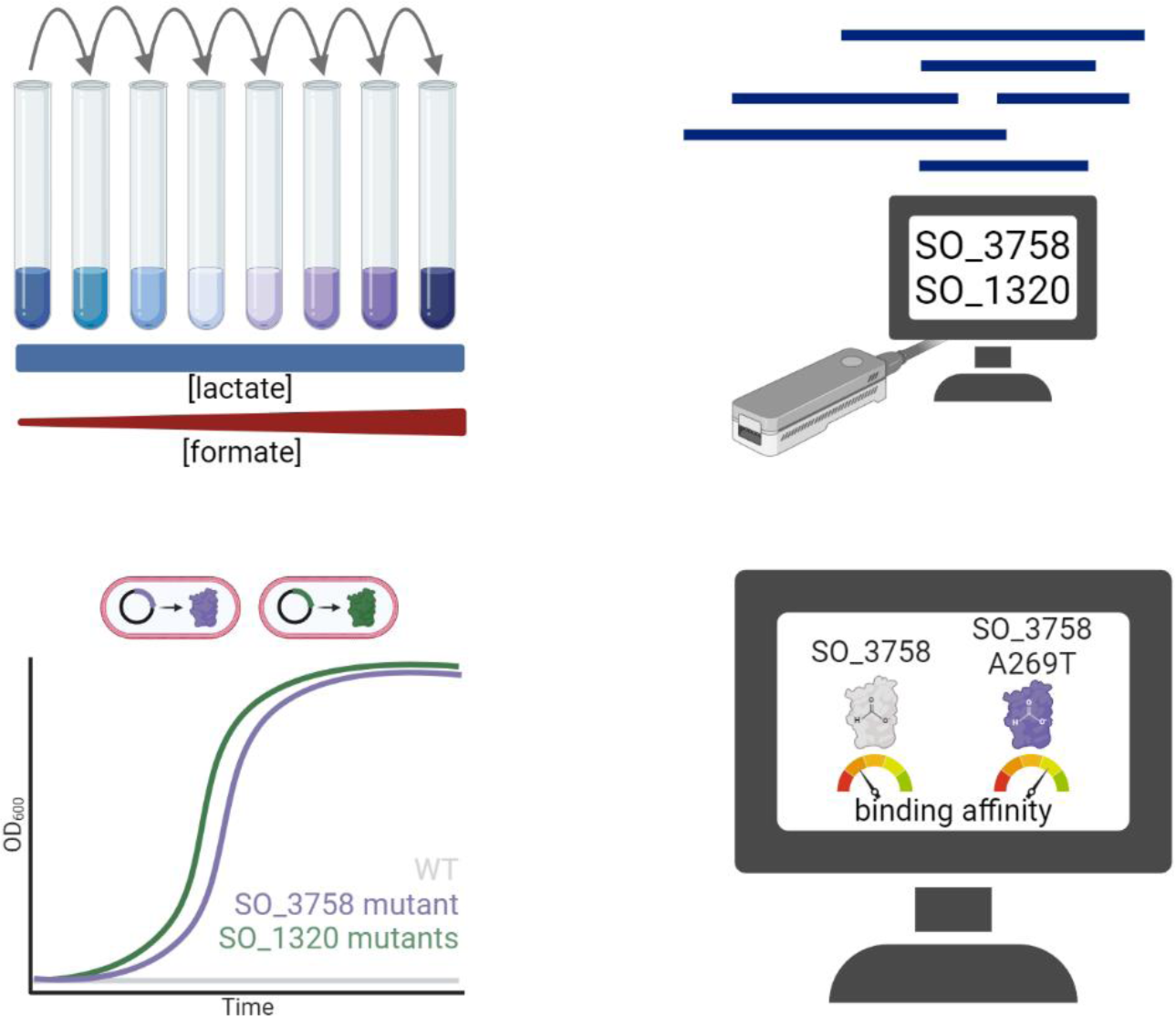

## Introduction

Microbial electrosynthesis (MES) is a promising technology that utilizes clean electricity and CO_2_ for the green production of biofuels, bioplastics, and platform chemicals (Dessì et al., 2021; Jensen et al., 2010; Jourdin & Burdyny, 2021). MES is mediated by electrochemically active organisms, which can utilize reducing power from an electrode and carbon from CO_2_ to generate target molecules through native or engineered metabolic reactions. Three different avenues of MES have been explored to date, each with their own limitations. The first method employs pure cultures of acetogens. At first glance, acetogens are an attractive MES catalyst, because they are autotrophic and some are electrochemically active; however, they suffer from a limited product scope, with acetate being the most abundant product (Prévoteau et al., 2020; Rosenbaum & Henrich, 2014; Schiel-Bengelsdorf & Dürre, 2012). Expansion of the product scope is hindered by the small synthetic biology toolbox for acetogens and their slow rate of electron uptake (Katsyv & Müller, 2020; Rosenbaum & Henrich, 2014). Another limitation of acetogens is the narrow understanding of their extracellular electron transfer (EET) mechanism, which makes it difficult to engineer improvements. At least five mechanisms have been proposed for EET in acetogens, but experimental evidence supporting any of these mechanisms is lacking, and it is not known whether a single EET mechanism is shared by all electrochemically active acetogens (Philips, 2020).

Some of these issues are addressed by the second major approach to MES, replacing the acetogen biocatalyst with mixed cultures. Mixed cultures have more success in producing a wider variety of products, although acetate production remains high (Arends et al., 2017; Ganigué et al., 2015; Raes et al., 2017). However, utilizing mixed cultures for MES is plagued by off-target methane production due to rising methanogen populations in the reactors over time (Prévoteau et al., 2020). A third approach uses model organisms in conjunction with exogenous electron carriers for MES. Model organisms like *E. coli* are more versatile because of their well understood metabolism and abundance of synthetic biology techniques (Harrington et al., 2015). However, scaling up MES reactions which require exogenous electron carriers is likely not feasible due to their cost (Harnisch et al., 2024; Rout et al., 2020). To make MES commercially competitive, a better microbial catalyst with well-understood EET mechanisms, metabolism, and synthetic biology tools is needed. One promising candidate is the electroactive bacterium *Shewanella oneidensis*. The discovery of the inward electron transfer pathway in this organism suggests that if coupled to a C1 assimilation pathway, *S. oneidensis* would likely serve as an improved MES biocatalyst (Ross et al., 2011; Rowe et al., 2018; Tefft & TerAvest, 2019).

*S. oneidensis* is a heterotrophic facultative anaerobe with the ability to utilize a wide variety of terminal electron acceptors for respiration including electrodes (Brutinel & Gralnick, 2012; Kouzuma, 2021). Extracellular electron transfer is made possible by the Mtr pathway, which includes the Mtr complex (MtrCAB), CctA, FccA, and CymA (Richardson et al., 2012). Quinone-linked oxidoreductases such as NADH dehydrogenase, formate dehydrogenase, and lactate dehydrogenase pass electrons obtained from their respective oxidation reactions to inner membrane quinones (Beblawy et al., 2018; Kane et al., 2016; Madsen & TerAvest, 2019). CymA, an inner membrane tetraheme cytochrome *c*, receives electrons from the quinone pool and subsequently passes them to periplasmic *c*-type cytochromes FccA and CctA to traverse the periplasm (Marritt et al., 2012; Myers & Myers, 1997; Sturm et al., 2015). On the other side of the periplasm, the cytochromes transfer the electrons to the outer membrane Mtr complex (MtrCAB) which transports the electrons to the outer surface of the cell (Richardson et al., 2012). The Mtr complex is composed of a porin (MtrB) which spans the outer membrane and encases two cytochromes *c*, one that interfaces with the periplasm (MtrA) and one that interfaces with the extracellular space (MtrC) (Coursolle et al., 2010; Coursolle & Gralnick, 2012). Once outside the cell, electrons can either directly reduce extracellular electron acceptors or be shuttled via extracellular flavins (Brutinel & Gralnick, 2012; Kouzuma, 2021).

Its well-studied, reversible Mtr pathway makes *S. oneidensis* a promising host for MES because it can efficiently exchange electrons with an electrode; however, this organism cannot fix CO_2_. *S. oneidensis* requires engineered CO_2_ fixation for its full potential as an effective MES biocatalyst to be realized. There are multiple potential methods for CO_2_ fixation, with two possibilities for initial conversions: carboxylation, such as in the Calvin cycle and the reductive tricarboxylic acid cycle, and reduction, such as in the Wood-Ljungdahl pathway and the reductive glycine pathway (Bar-Even et al., 2012; Cotton et al., 2018; François et al., 2020). Pathways which first reduce CO_2_ to formate have been found to support higher biomass yields due to utilizing ATP more efficiently (Bar-Even et al., 2012; Cotton et al., 2018). Because of this, engineering CO_2_-fixation in *S. oneidensis* via a reduction-first pathway would be a promising strategy for generating an MES-compatible host. CO_2_ can be reduced to formate via reversible formate dehydrogenases (Amao, 2018; Calzadiaz-Ramirez & Meyer, 2022) and formate can be assimilated through a host of methods such as the reductive glycine pathway, the serine cycle, and reversed activity of pyruvate-formate lyase (Bar-Even, 2016). Additionally, formate-assimilating *S. oneidensis* could also be utilized for MES when an electrode with adsorbed formate dehydrogenase is supplied, as such electrodes have been shown to freely interconvert CO_2_ and formate with low overpotentials (Amao, 2018; Reda et al., 2008). *S. oneidensis* is not naturally tolerant of very high formate concentrations, necessitating greater tolerance to be acquired if a CO_2_-reducing electrode is used or a reduction-first CO_2_ fixation pathway is expressed. In this work, we utilized adaptive laboratory evolution to create formate-tolerant strains of *S. oneidensis* as a first step toward developing an MES-compatible host. We found two genes that are responsible for increasing formate tolerance in this organism and explore their application toward a future CO_2_ fixing strain of *S. oneidensis*. We also show that transfer of one formate tolerance mechanism to another host, *Zymomonas mobilis*, is possible, demonstrating that conferral of formate tolerance to other species is possible through the expression of one gene.

## Methods

### Strains and Plasmids

Strains and plasmids used in this study are listed in **Table S1**. Plasmids for mutant gene overexpression were prepared by cloning the WT or mutant gene from WT or mutant genomic DNA via PCR with primers (**Table S2**) that added regions homologous to the backbone vector pRL814 (Ghosh et al., 2019) at the NdeI and HindIII restriction sites. WT copies of SO_3758 and SO_1320 were amplified from WT *S. oneidensis*, SO_3758_A269T was amplified from MGC002, SO_1320_V106I was amplified from MGC007, SO_1320_V106F was amplified from MGC003, and SO_1320_S52I was amplified from MGC008. Amplified genes were purified via the Wizard PCR clean- up kit (Promega). The pRL814 (containing green fluorescent protein (GFP) under control of *P*_T7A1-O34_) backbone was digested with NdeI and HindIII-HF (NEB). The digested backbone and amplified genes were ligated using NEBuilder HiFi DNA Assembly Master Mix (NEB) and transformed into chemically competent *E. coli* WM3064, a 2,6-diaminopimelic acid (DAP) auxotroph. Plasmids were transferred to WT *S. oneidensis* MR-1 via conjugal transfer using *E. coli* donor WM3064 (Coursolle & Gralnick, 2012) or into *Z. mobilis* ZMDR (Lal et al., 2021) using *E. coli* donor WM6026 as described previously (Felczak et al., 2021; Lal et al., 2019). Mobile CRISPRi knockdown vectors were prepared from the backbone pJMP2846 as described previously, with sgRNAs targeting *yadS* (Ford et al., 2022; Peters et al., 2019). The CRISPRi vectors are kanamycin resistant and encode dCas9 and the sgRNA. The assembled CRISPRi vectors were transformed into chemically competent *E. coli* WM3064. The CRISPRi vectors were transferred to WT *S. oneidensis* via Tn7-based triparental conjugation using donor strains *E. coli* WM3064 with the CRISPRi vector and transposase-containing *E. coli* WM3064 pJMP2644 as described previously (Peters et al., 2019). After the mating period, counter-selection was performed on LB agar plates supplemented with 50 μg ml^-1^ of kanamycin and without DAP. Single *S. oneidensis* colonies were screened for the correct sgRNAs by PCR using the sgRNA_check and appropriate sgRNA reverse primers (**Table S2**).

### Culturing

*S. oneidensis* strains were pre-cultured in 2 ml LB with the appropriate antibiotic overnight at 30 °C with shaking at 250 rpm. The pre-cultures were washed three times and resuspended in basal M5 -caa medium (1.29 mM K2HPO_4_, 1.65 mM KH_2_PO_4_, 7.87 mM NaCl, 1.70 mM NH_4_SO_4_, 475 μM MgSO_4_·7H_2_O, 10 mM HEPES, pH 7.2). Minimal lactate medium was used for all experimental *S. oneidensis* cultures unless otherwise stated and was comprised of basal M5 -caa, supplemented with Wolfe’s mineral solution, Wolfe’s vitamin solution without riboflavin, and 20 mM D,L-lactate. Formate and isopropyl β-D-1-thiogalactopyranoside (IPTG) were added to the appropriate cultures at the specified concentrations. Washed pre-cultures were used to inoculate the medium to an initial OD_600_ of 0.02. Cultivation took place at 30 °C with shaking at 250-275 rpm. Culture volumes are specified for each experiment.

*Z. mobilis* strains were pre-cultured overnight in ZRMG liquid medium (1% yeast extract, 2% glucose, 15 mM KH_2_PO_4,_ and 100 μg/ml spectinomycin) at 30°C, with shaking at 250 rpm. Formate tolerance growth experiments were conducted in ZRMG medium and were inoculated to an initial OD_600_ of 0.01 from the pre-culture. Formate was added at the specified concentrations. Growth of *Z. mobilis* strains were conducted with basal expression of GFP, SO_3758, or SO_3758_A269T from pRL814. Cultivation occurred in 200 μl of medium at 30 °C with shaking at 250-275 rpm. Growth of *S. oneidensis* and *Z. mobilis* strains in 200 μl cultures were monitored on a H1M BioTek Plate Reader.

### Evolution of formate tolerance and genome sequencing

Formate tolerance in *S. oneidnesis* was evolved by continuously subculturing the parent strains (WT, JG2957 JG2955) in 2 ml minimal lactate medium with increasing concentrations of formate. Pre-cultures of the parent strains were prepared as described above and were used to inoculate three 2 ml cultures of minimal lactate medium with 1 mM formate to an initial OD_600_ of 0.02. The three or four cultures for each parent strain represented a separate evolution line: line A, line B, line C, and line D. Cultures were incubated at 30 °C with shaking at 250 rpm. 2 μl aliquots were taken daily to monitor OD_600_ using an Eppendorf G1.0 microcuvette and an Eppendorf BioSpectrometer. When cultures reached an OD_600_ of 0.2 or higher, they were subcultured into a fresh 2 ml culture with minimal lactate medium and 5 mM formate. Strains were continuously cultured in this way with the formate concentration increasing in each subculture. The formate concentration progression was: 1 mM, 5 mM, 10 mM, 20 mM, 30 mM, 40 mM, 50 mM, 60 mM, 70 mM, 80 mM, 90 mM, 100 mM. Strains spent only one round of culturing in each formate concentration before moving to the next highest concentration. Cryogenic stocks were prepared from each of the populations when the 100 mM cultures reached an OD_600_ of 0.2 or higher. Cryogenic stocks from each evolution line were streaked onto LB agar plates, and multiple single colonies from each evolution line were saved as pure cryogenic stocks. Genome extraction, library preparation, and Oxford Nanopore sequencing was performed by Plasmidsaurus.

### Bioinformatics

Raw genome sequencing reads were trimmed using Porechop v0.2.4 (https://github.com/rrwick/Porechop.git) to remove Nanopore sequencing adapters. Trimmed sequencing reads were aligned to the *S. oneidensis* MR-1 reference genome (Accession: NC_004347.2) and reference megaplasmid (Accession: NC_004349.1) and analyzed for mutations using Breseq v0.38.0 (Deatherage & Barrick, 2014). Deviations of the evolved strains from the reference sequences were compared to the deviations of the parent strains from the reference sequences. Deviations which appeared in the evolved strains but not in the parent strains were compiled in **Table S3** and classified as mutations. Mutations which appeared in multiple evolved strains were of particular interest. A sodium-dependent bicarbonate transporter (SO_3758) and a DUF2721-containing protein (SO_1320) were experimentally investigated for utility in formate tolerance.

iPro54-PseKNC and Sigma70Pred were used to search for *yadS* σ^54^ and σ^70^ promoters at the end of the SO_1320. An 81-nucleotide long sequence containing the end of SO_1320 and the first eight nucleotides of *yadS* were used as the query. (Lin et al., 2014; Patiyal et al., 2022).

Amino acid sequences of WT and mutant versions of SO_3758 were aligned to the sodium-dependent bicarbonate transporter SbtA from cyanobacteria *Synechocystis* sp. PCC 6803 substr. Kazusa (Accession: P73953) using T-coffee simple multiple sequence alignment(Notredame et al., 2000).

Tools DeepGO and DeepTMHMM were used to analyze the amino acid sequence of SO_1320 (Hallgren et al., 2022; Kulmanov et al., 2018). AlphaFold v2.3.2 was used to predict structures of SO_3758_A269T, SO_1320_V106I, SO_1320_V106F, and SO_1320_S52I (Jumper et al., 2021).

Predicted structures for SO_3758 (Accession: Q8EAY3) and SO_1320 (Accession: Q8EHA9) were downloaded from the AlphaFold database (https://alphafold.com/). Ligand binding prediction was performed with AutoDock Vina v1.1.2 (Trott & Olson, 2010). PyMOL v2.5.8 was used for visualization of protein and ligand models, identifying polar interactions between the ligand and surrounding amino acids, and calculating distances between interacting atoms (Schrödinger LLC, 2015).

## Results

Two *S. oneidensis* strains were used in the testing and evolution of formate tolerance: wild-type (WT) and JG2957. *S. oneidensis* strain JG2957 has three major FDHs deleted (*fdhA1B1C1*, *fdhA2B2C2*, and *fdnGHI*) and was included to prevent oxidation of formate to CO_2_ and provide a deeper understanding of formate tolerance.

First, to assess the native formate tolerance of *S. oneidensis*, WT and JG2957 were cultivated in 200 μl minimal lactate medium with varying concentrations of formate (**Figure 1**). Both strains had the shortest lag phase in medium without formate. With even as little as 1 mM formate, growth was impacted. With increasing formate concentrations, the strains’ lag phase also increased. Both strains were only capable of tolerating up to 10 mM formate. The low formate tolerance in these strains could present an issue in a *S. oneidensis* strain which has CO_2_ reductase activity.

**Figure 1.**
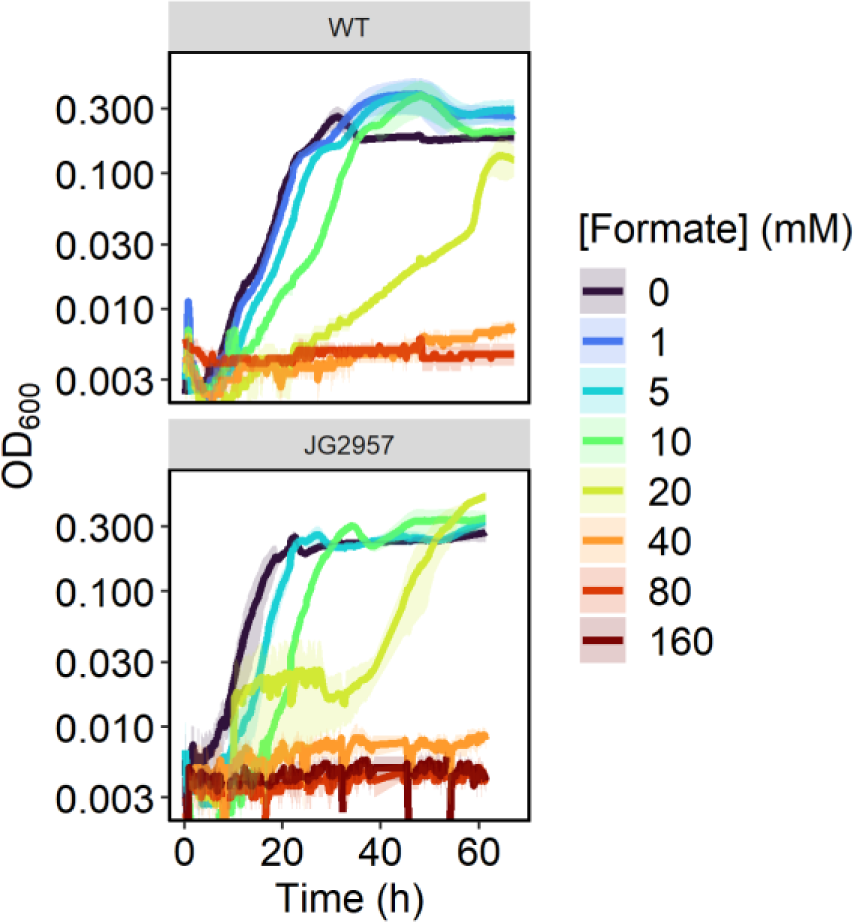
Growth curves of MR-1 and JG2957 in 200 μl minimal medium with 20 mM lactate and varying concentrations of formate.

Formate toxicity can be a result of cytochrome *c* oxidase inhibition and/or reduction of proton motive force due to cytoplasm acidification from the protonated acid (Cotton et al., 2020; Nicholls, 1975; Warnecke & Gill, 2005). Because these issues are not easily solved with a genetic engineering approach, we opted to generate a formate-tolerant strain of *S. oneidensis* through adaptive laboratory evolution. This was accomplished through continuous subculturing of the parent strains (WT and JG2957) in minimal lactate medium while increasing the formate concentration in the medium for each subculture, with a goal of improving tolerance to 100 mM formate. Multiple separate evolutions were conducted for each parent strain. Surprisingly, tolerance to 100 mM formate evolved within 30 days in seven evolution lines (**Figure S1**), with the subculturing frequency increasing in the later stages of the evolution, indicating acquired mutation(s) assisting with formate tolerance (**Figure 2**).

**Figure 2.**
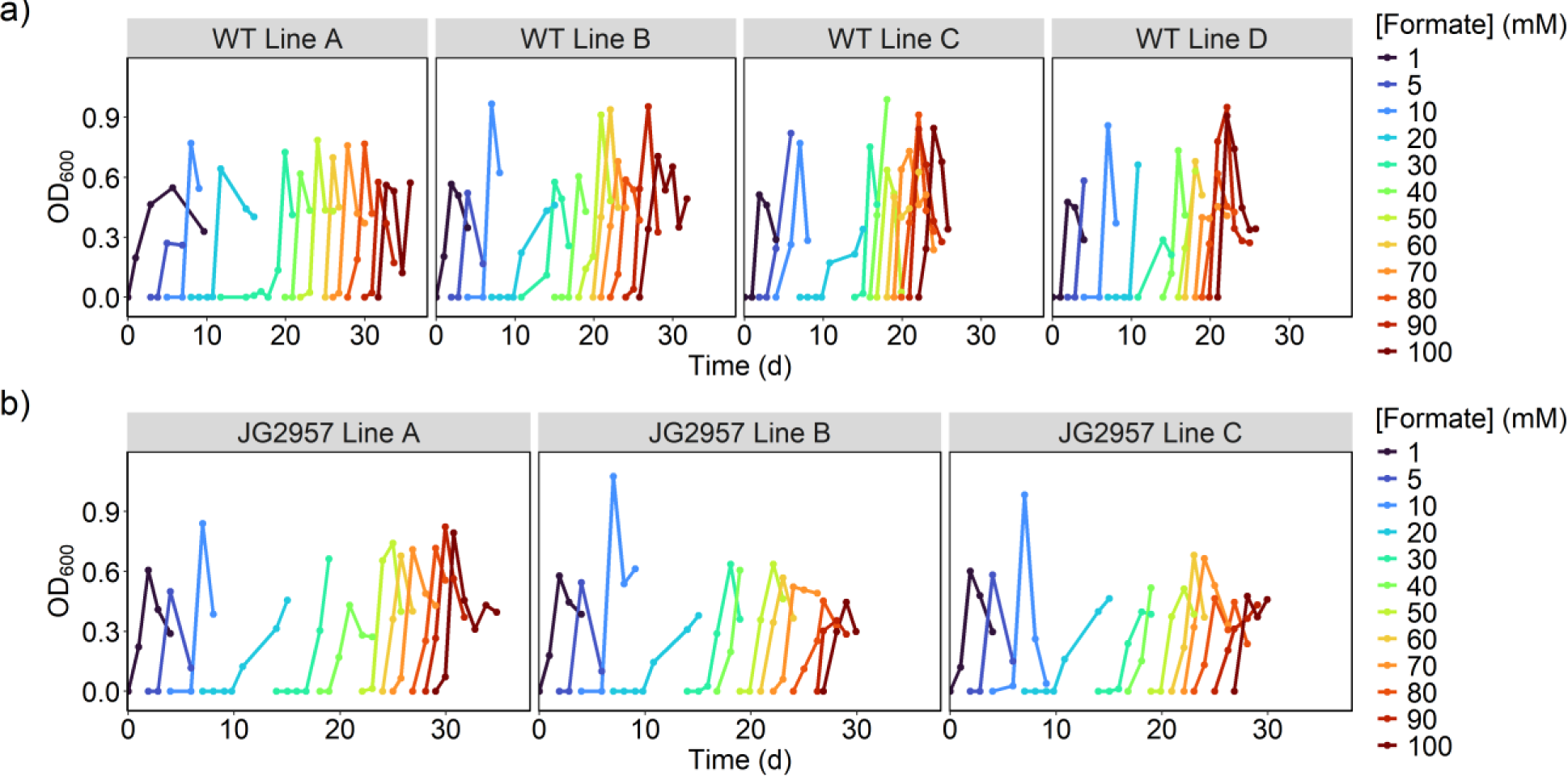
Growth curves of directed evolution subculturing in increasing formate concentrations of parent strains a) WT and b) JG2957. Each panel displays growth curves of a replicate evolution line. Evolutions were ceased after formate concentration reached 100 mM.

Single colonies from the seven evolution lines capable of growth on 100 mM formate were isolated. Their formate tolerance was compared to their associated parent strain by culturing in 200 μl minimal lactate media with 100 mM formate in triplicate (**Figure 3**). Both parent strains were incapable of growth on lactate when supplemented with 100 mM formate, while almost all the isolated evolved colonies were capable of growth, although not without a long lag phase of 60-90 hours. The evolutions from WT resulted in four lines (A, B, C, and D) which showed different growth phenotypes with lines A, C, and D having a shorter lag phase than line B. Less clear patterns emerge from strains evolved from JG2957. Two of the line A colonies and one line B colony could grow the most optimally under the conditions tested, while the other colonies from those lines more closely resembled the growth patterns of colonies from line C.

**Figure 3.**
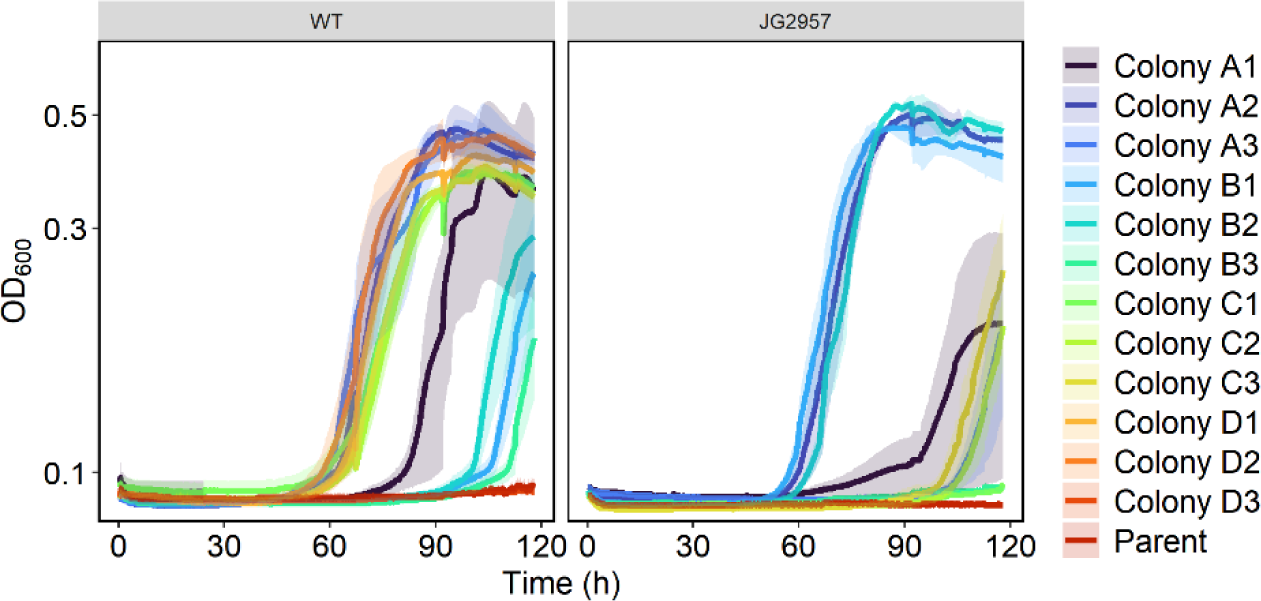
Growth curves of single colonies in triplicate isolated from evolution lines at 100 mM formate compared to their associated parent strain. Strains were cultured in 200 μl of minimal lactate medium with 100 mM formate.

One colony from each line was selected for further analysis. These seven new formate- tolerant *S. oneidensis* strains were renamed MGC001 through MGC007 (**Table S1**). Each formate- tolerant strain and the parent strains were cultivated in 50 ml minimal lactate medium with 100 mM formate, and metabolites were analyzed via HPLC (**Figure 4**, **Figure S2**). Cultures were inoculated to a starting OD_600_ five times greater than previous cultures to allow the parent strains to grow under the high-formate conditions. This provided a means for comparison between the evolved and parent strains’ metabolite concentrations (**Figure S2**). Even with the increased inoculum volume, WT and JG2957 had a longer lag phase than the evolved formate- tolerant strains. The evolved formate-tolerant strains did not have as long of a lag phase as when inoculated to a lower

**Figure 4.**
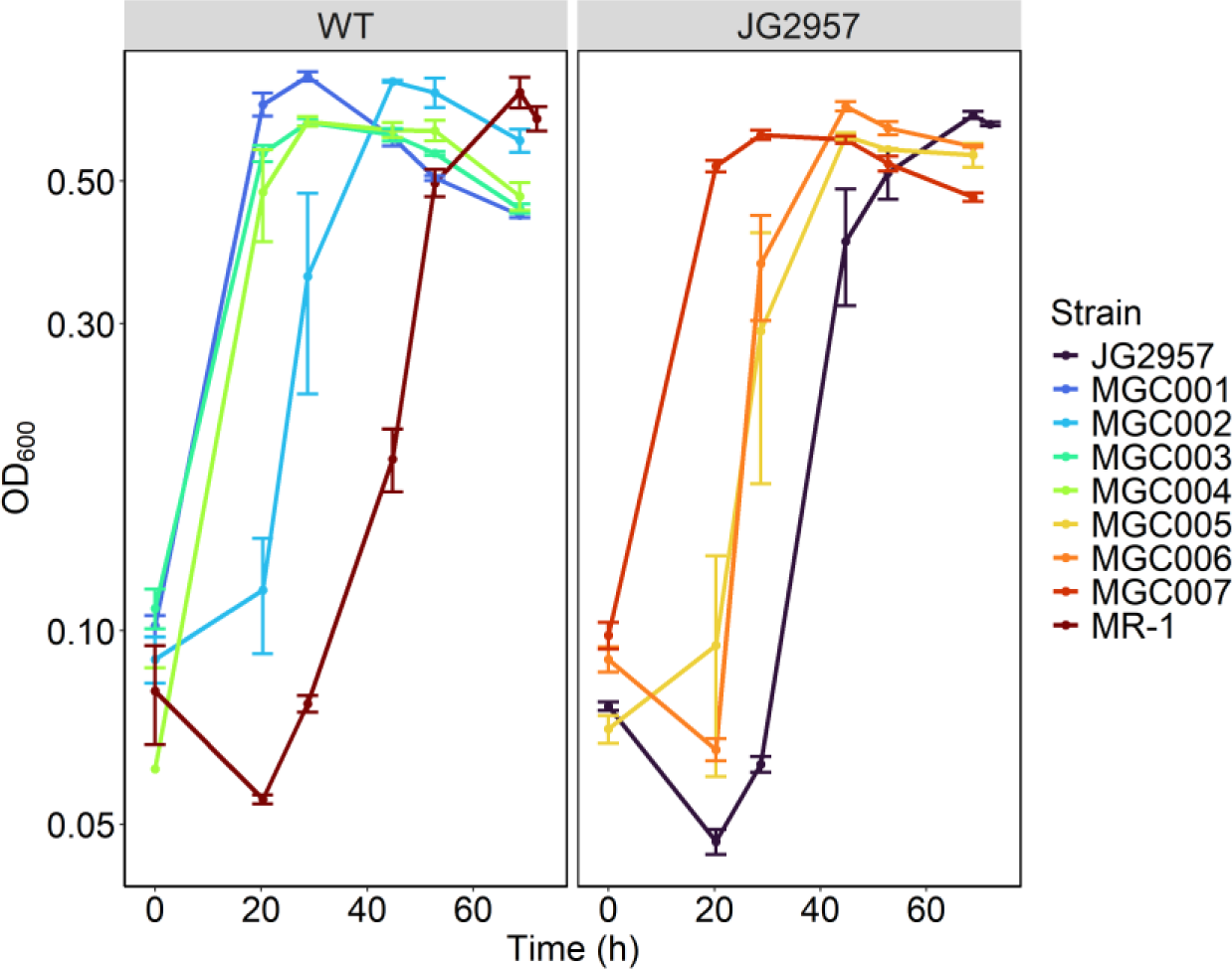
Growth curves of formate-tolerant strains cultivated in 50 ml minimal lactate medium with and 100 mM formate.

OD600. All provided lactate was consumed in all strains (**Figure S2**). Formate levels remained constant throughout cultivation of both the parent strains and the evolved formate tolerant strains, indicating that the evolved method of formate tolerance does not include formate oxidation or formate assimilation.

To determine the mutations that enabled increased formate tolerance, the genomes of the seven formate-tolerant strains and the two parent strains were sequenced. Raw sequencing reads were processed with porechop to remove sequencing adapters and breseq to align the reads to the reference sequences and identify mutations. Mutations that did not exist in the parent strain were flagged (**Table S3**). A transition mutation of G to A in SO_3758 (SO_RS17510), encoding the core domain of a Sbt-like sodium-dependent bicarbonate transporter, occurred in three of the seven formate-tolerant strains (MGC002, MGC005, and MGC006). This resulted in the amino acid sequence mutation A269T, changing a nonpolar, hydrophobic residue to a bulkier polar, hydrophilic residue. Considering that formate and bicarbonate have very similar structures (**Figure 5**), we hypothesized two potential effects of this mutation. First, the mutation could have provided greater specificity toward formate and provided SO_3758 with formate efflux activity, preventing toxic intracellular accumulation. Second, the mutation may have decreased formate specificity and prevented SO_3758 from importing toxic levels of formate.

**Figure 5.**
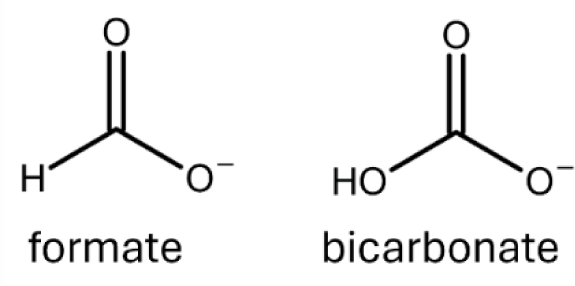
Chemical structures of formate and bicarbonate.

In three of the remaining formate-tolerant strains (MGC003, MGC004, and MGC007), a mutation in SO_1320 (SO_RS06120), a protein of unknown function, was observed. MGC007 had a G to A transition mutation resulting in a V106I amino acid sequence change and MGC003 and MGC004 had a G to T transversion mutation resulting in a V106F amino acid sequence change. We also carried out preliminary adaptive laboratory evolution on using *S. oneidensis* parent JG2955 which has two of the three formate dehydrogenases deleted (*fdhA1B1C1* and *fdhA2B2C2*). Interestingly, one strain (MGC008) isolated from this evolution also had a mutation in SO_1320 but in residue 52 where a serine was converted to an isoleucine. The two residue 106 mutations resulted in a bulkier amino acid in that position, and the S52I mutation replaced a polar, hydrophilic residue with a nonpolar, hydrophobic residue.

SO_1320 is upstream of SO_1319 (*yadS*) and SO_1318. To determine whether the mutations in SO_1320 affected the promoter sequence of downstream *yadS*, an 81 bp unmutated nucleotide sequence containing the end of SO_1320 and first eight nucleotides of *yadS* were analyzed by iPro54- PseKNC and Sigma70Pred to search for σ^54^ and σ^70^ promoters (Lin et al., 2014; Patiyal et al., 2022). These analyses revealed no σ^54^ promoter in this location, but Sigma70Pred predicted the region contained a σ^70^ promoter. The same 81-nucleotide sequence including the G to A or G to T mutations resulting in V106 change to phenylalanine or isoleucine in the formate-tolerant strains were also analyzed by Sigma70Pred. In both cases, the sequence is still predicted to contain a σ^70^ promoter, suggesting that the mutations do not alter the transcription of downstream *yadS*. This finding, along with the observation that a mutation elsewhere in SO_1320 also results in formate tolerance, indicates that the mutations in SO_1320 alter this protein’s activity in a way that helps the strains withstand greater formate concentrations.

To confirm that the SO_1320 mutations do not impact the transcription of *yadS*, two *S. oneidensis* IPTG-inducible CRISPRi knockdown strains were prepared, each targeting *yadS*. The *yadS* knockdown strains and the non-targeting control were cultivated in minimal lactate medium with 20 mM formate and varying levels of induction. If decreased expression of *yadS* was responsible for the improvement of formate tolerance, the *yadS* CRISPRi targets were expected to show increased formate tolerance with greater induction levels. However, the growth curves reveal that knockdown of *yadS* does not improve *S. oneidensis* formate tolerance over the non-targeting strain, verifying that the mutations in SO_1320 do not affect *yadS* expression (**Figure S3**).

To confirm that mutations observed in SO_3758 and SO_1320 were responsible for enhanced formate tolerance, we attempted to confer formate tolerance to WT *S. oneidensis* by overexpressing the mutant sequences. Mutant genes were cloned from the formate-tolerant strains and placed on vector pRL814 under control of an IPTG-inducible promoter. The vectors carrying the mutants were transformed into *E. coli* and subsequently transferred to WT *S. oneidensis* via conjugation. The *S. oneidensis* strains harboring SO_3758, SO_3758_A269T, SO_1320, SO_1320_V106I, SO_1320_V106F, or SO_1320_S52I were cultivated in 200 μl of minimal lactate medium with varying formate concentrations and the WT or mutant gene was overexpressed with 100 μM IPTG (**Figure 6**). All strains, whether expressing WT or mutant versions of the genes were capable of growth without formate, showing that overexpression of these proteins was not toxic. The strains expressing the WT versions of SO_3758 and SO_1320 were only able to grow robustly in the no formate conditions. The strains expressing any of the mutant sequences grew in elevated formate levels up to 40 mM with a final OD_600_ similar to growth in medium without formate. None of the strains tolerated 80 mM formate, indicating that other mutations accumulated during evolution of the formate-tolerant strains could enhance their ability to withstand elevated levels of formate. This analysis confirmed that the identified mutations in SO_3758 and SO_1320 improved formate tolerance of WT *S. oneidensis*.

**Figure 6.**
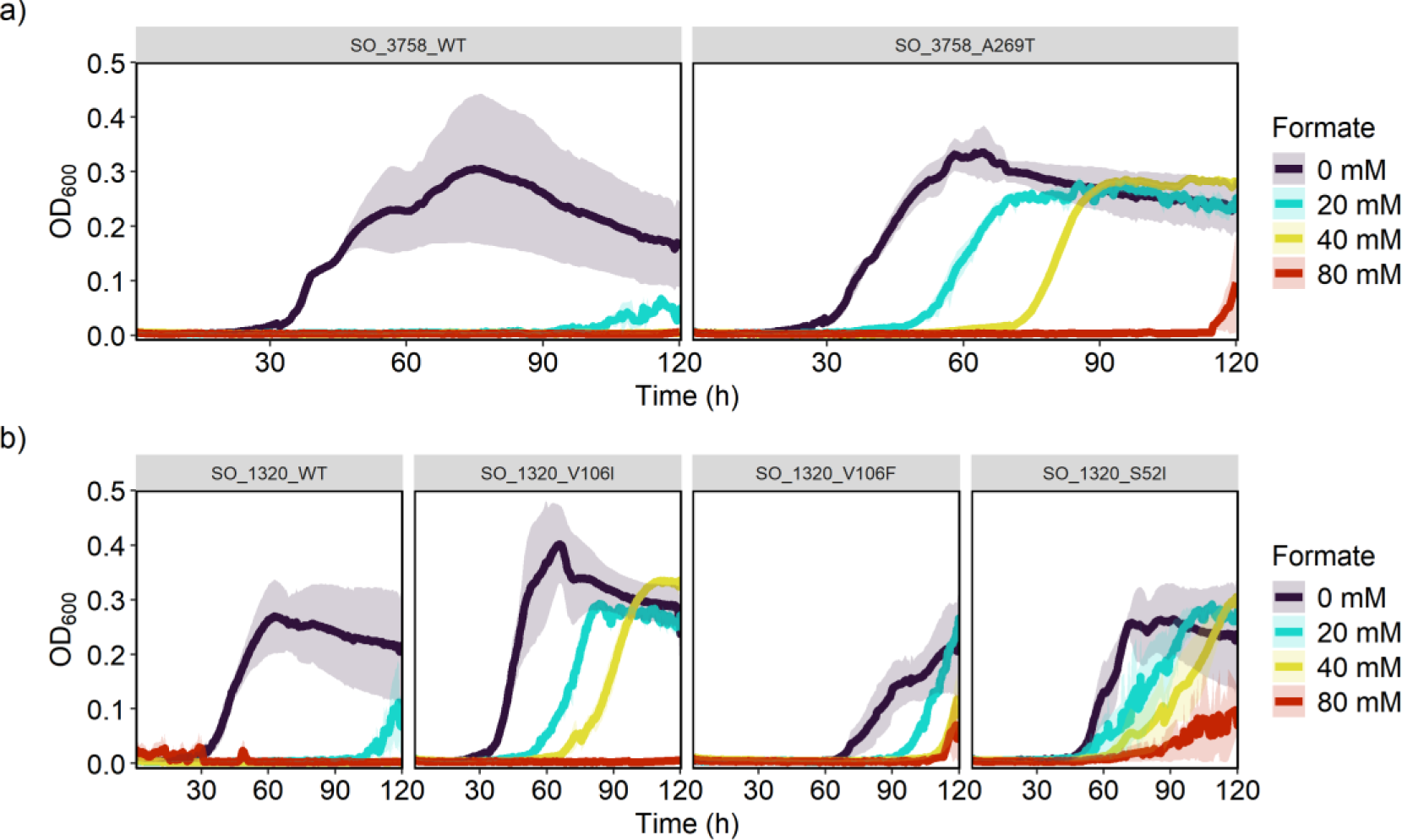
Growth curves of WT *S. oneidensis* harboring an IPTG-inducible expression vector containing WT or mutant versions of a) SO_3758 or b) SO_1320 in 200 μl minimal lactate medium with 100 µM IPTG and varying formate concentrations.

To predict the effect of the A269T mutation in the transporter SO_3758, we modeled the binding of bicarbonate and formate to the WT and mutant versions of the protein. To reveal the binding site for modeling substrate binding, the *S. oneidensis* SO_3758 amino acid sequence was aligned with the only Sbt-like sodium-dependent bicarbonate transporter core domain with a solved structure (SbtA from cyanobacteria) using the multiple sequence alignment (MSA) tool T-Coffee (Liu et al., 2021; Notredame et al., 2000). The alignment revealed that A269 in SO_3758 corresponds to residue S323 in SbtA (**Figure S4**). While S323 has not been found to be directly involved in binding sodium or bicarbonate, it is the single residue that separates the two binding pockets, with S322 binding sodium and S324 binding bicarbonate (Fang et al., 2021; Liu et al., 2021). These substrate-binding serines are conserved in SO_3758, with SO_3758 S268 corresponding to SbtA S322 and SO_3758 S270 corresponding to SbtA S324 (**Figure S4**). The MSA revealed that SO_3758 residues V99, S100, S270, and I272 are likely part of the bicarbonate binding pocket.

Protein models of the wild-type and mutated versions of SO_3758 were generated using AlphaFold (Jumper et al., 2021), and the binding of bicarbonate and formate were predicted with Autodock Vina (Trott & Olson, 2010). The predicted local distance difference test (pLDDT) scores were calculated for each residue in the AlphaFold model and are depicted in **Figure 7**. The binding pocket residues were predicted to have pLDDT scores of high (90 > pLDDT > 70) to very high (pLDDT > 90) confidence. The docking grid was set to encompass residues predicted by MSA to be part of the binding pocket (V99, S100, S270, I272), A or T269, and nearby residues. The modeling of bicarbonate binding to SO_3758_WT predicts that it forms polar interactions with V99, A101, Y271, I272, and A273 in the predicted native binding site, referred to here as site 1 (**Figure 8a**). Bicarbonate binding to site 1 has a predicted binding affinity of -3.2 kcal mol^-1^ (**Table 1**). Formate is also predicted to bind to site 1 in SO_3758_WT, forming polar interactions with V99, S270, Y271, and I272 and having a lower predicted binding affinity of -2.3 kcal mol^-1^ (**Figure 8b**).

**Figure 7.**
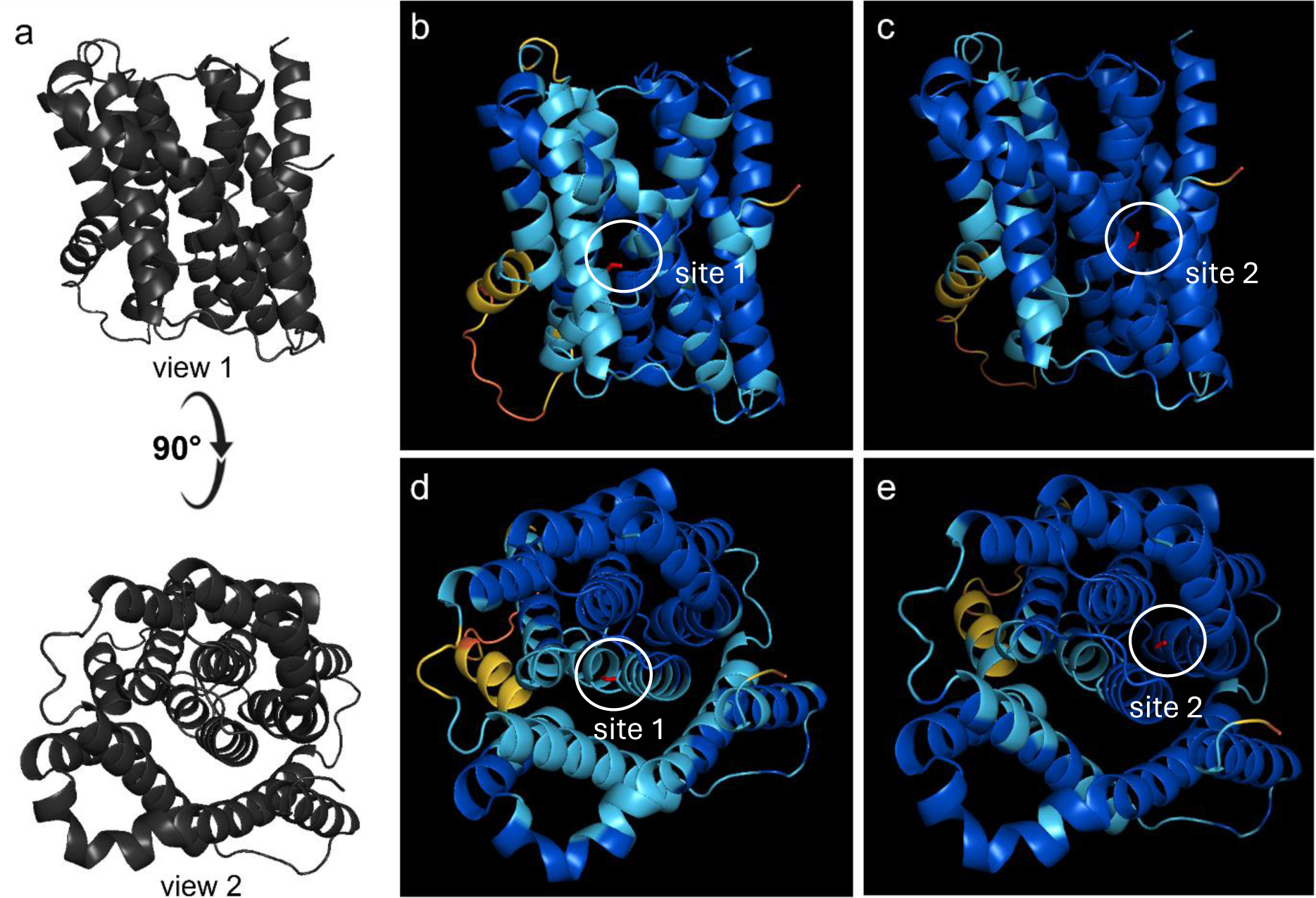
Representations of pLDDT scores from SO_3758 AlphaFold structural predictions. Residue color depicts pLDDT score. Dark blue represents a very high score (pLDDT > 90). Light blue represents a confident score (90 > pLDDT > 70). Yellow represents a low score (70 > pLDDT > 50). Orange represents a very low score (pLDDT < 50). a) General structure of SO_3758 shown at two angles. View 1 is shown in panels b and c. View 2 is shown in panels d and e. b) View 1 of WT SO_3758. Formate is colored red and shown binding to site 1. c) View 1 of SO_3758_A269T. Formate is colored red and shown binding to site 2. d) View 2 of WT SO_3758. Formate is colored red and shown binding to site 1. c) View 2 of SO_3758_A269T. Formate is colored red and shown binding to site 2.

**Figure 8.**
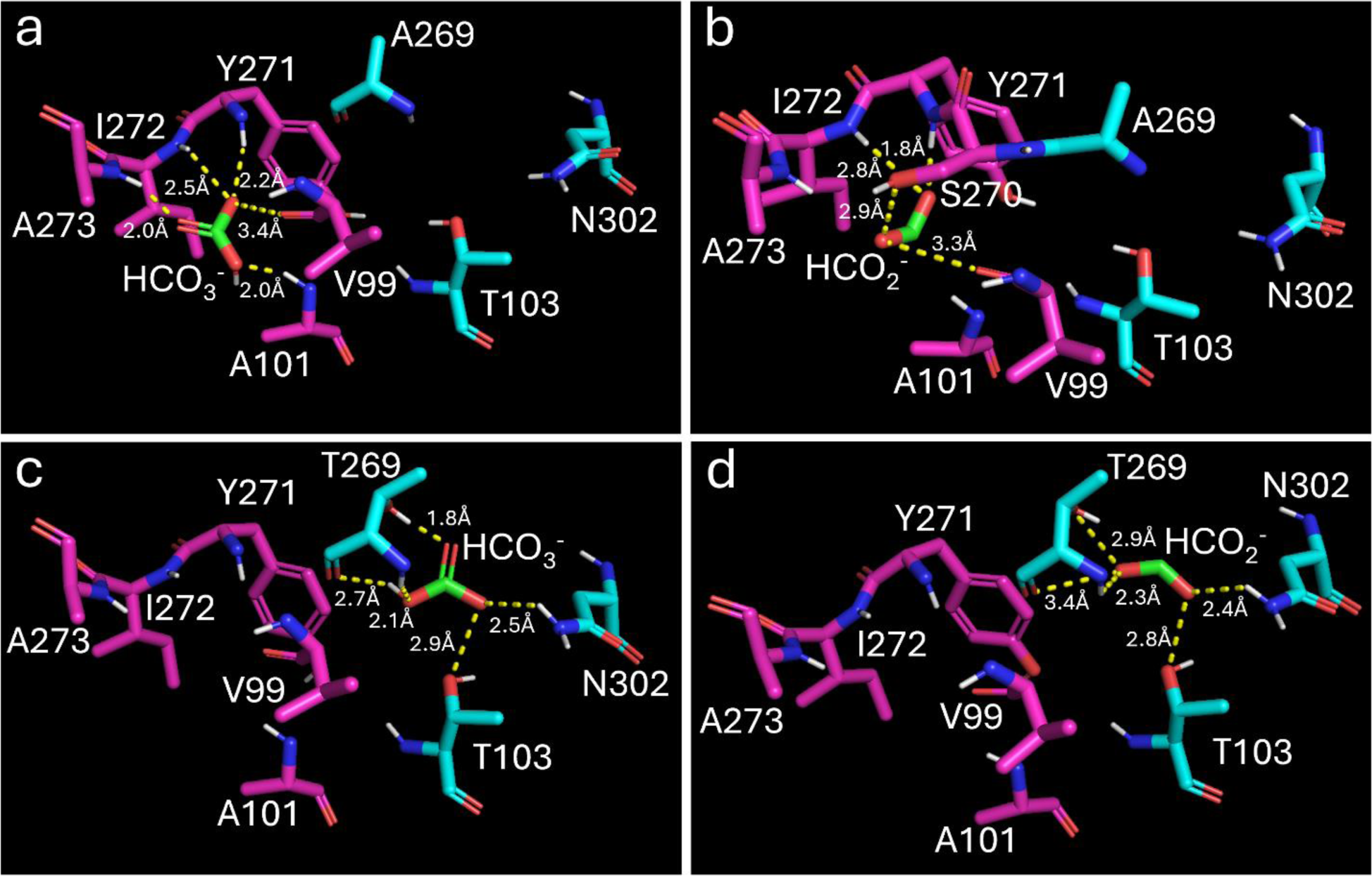
Modeled binding of bicarbonate (HCO_3-_) to a) SO_3758_WT and c) SO_3758_A269T, and of formate (HCO_2-_) to b) SO_3758_WT and d) SO_3758_A269T. Residues in the native bicarbonate binding site (site 1) are shown in pink and residues in site 2 are shown in cyan. Formate and bicarbonate are shown in green. Nitrogen atoms are colored blue, oxygen atoms are colored red, and hydrogen atoms are colored white. Polar interactions between bicarbonate or formate and interacting residues are shown as yellow dotted lines and are labeled with the distance between the interacting atoms. Both bicarbonate and formate form polar interactions with the mutated residue (T269) in SO_3758_A269T that contributes to the tighter binding of these substrates in the mutated protein.

**Table 1.** Predicted binding affinities of bicarbonate and formate to site 1 and site 2 of WT and mutant versions of SO_3758.

|  | Binding affinity to SO_3758_WT |  | Binding affinity to SO_3758_A269T |  |
| --- | --- | --- | --- | --- |
|  | Site 1(kcal mol<sup>-1</sup>) | Site 2 (kcal mol<sup>-1</sup>) | Site 1 (kcal mol<sup>-1</sup>) | Site 2 (kcal mol<sup>-1</sup>) |
| Bicarbonate | -3.2 | -2.9 | -3.4 | -3.7 |
| Formate | -2.3 | NA | -2.9 | -3.1 |

When modeled with SO_3758_A269T, bicarbonate is still predicted to bind to site 1 and form polar interactions with the same residues with a slightly higher binding affinity of -3.4 kcal mol^-1^. However, the greatest predicted binding affinity of -3.7 kcal mol^-1^ is in a binding pocket with the mutated T269 residue, as well as residues T103 and N302, here referred to as site 2 (**Figure 8c**). This indicates that the A269T mutation altered the native bicarbonate binding site 1 slightly to promote tighter binding of the native substrate, even though residue 269 is not itself a part of site 1. More interestingly, it shows that the mutation in residue 269 alters site 2 to increase affinity for the substrate. Finally, when formate binding is modeled with SO_3758_A269T, the greatest predicted binding affinity (-3.1 kcal mol^-1^) is also in site 2 with T103, N302, and the mutated residue T269 (**Figure 8d**). This brings the predicted binding affinity of formate much closer to that of bicarbonate in site 1 of the native SO_3758 (-3.2 kcal mol^-1^), suggesting that formate is a suitable substrate for this mutant. The predicted interaction of formate with threonine 269 in SO_3758_A269T but not with alanine in SO_3758_WT also suggests that this mutation was beneficial in promoting formate binding.

The two binding sites are predicted to bind both substrates in SO_3758_A269T, but formate is not predicted to bind to site 2 in WT SO_3758 (**Table 1**). Site 1 binds both bicarbonate and formate better than site 2. The opposite is true for SO_3758_A269T where site 2 with residue threonine 269 binds both substrates better and increases the predicted binding affinity of formate to a similar level that is predicted for bicarbonate binding in the native protein. Based on this modeling, we anticipated that the A269T mutation in SO_3758 provided MGC002, MGC005, and MGC006 with greater formate efflux capacity to prevent the toxic accumulation of formate inside the cells.

To determine whether SO_3758_A269T could also confer formate tolerance to other species, we expressed SO_3758_WT and SO_3758_A269T in *Zymomonas mobilis*. This organism natively has greater formate tolerance than *S. oneidensis* and is a native ethanol producer, making it a promising bio-production platform (Braga et al., 2021; He et al., 2014). To determine whether SO_3758_A269T could improve formate tolerance in *Z. mobilis*, the organism was grown in ZRMG medium with various formate concentrations either with the control vector (pRL814 expressing GFP), pRL814-SO_3758, or pRL814-SO_3758_A269T (**Figure 9**). With its greater native formate tolerance, *Z. mobilis* carrying the control vector was able to maintain similar growth in 20 mM and 60 mM formate as without formate. However, the control vector strain growth was impacted at 80 mM formate. The basal expression of either the WT or mutant versions of SO_3758 allowed *Z. mobilis* to maintain similar growth across all formate concentrations, indicating that both SO_3758_WT and SO_3758_A269T ability to improve formate tolerance in *Z. mobilis*. Inducing expression of either the WT or mutant versions of SO_3758 was toxic for *Z. mobilis* (data not shown).

**Figure 9.**
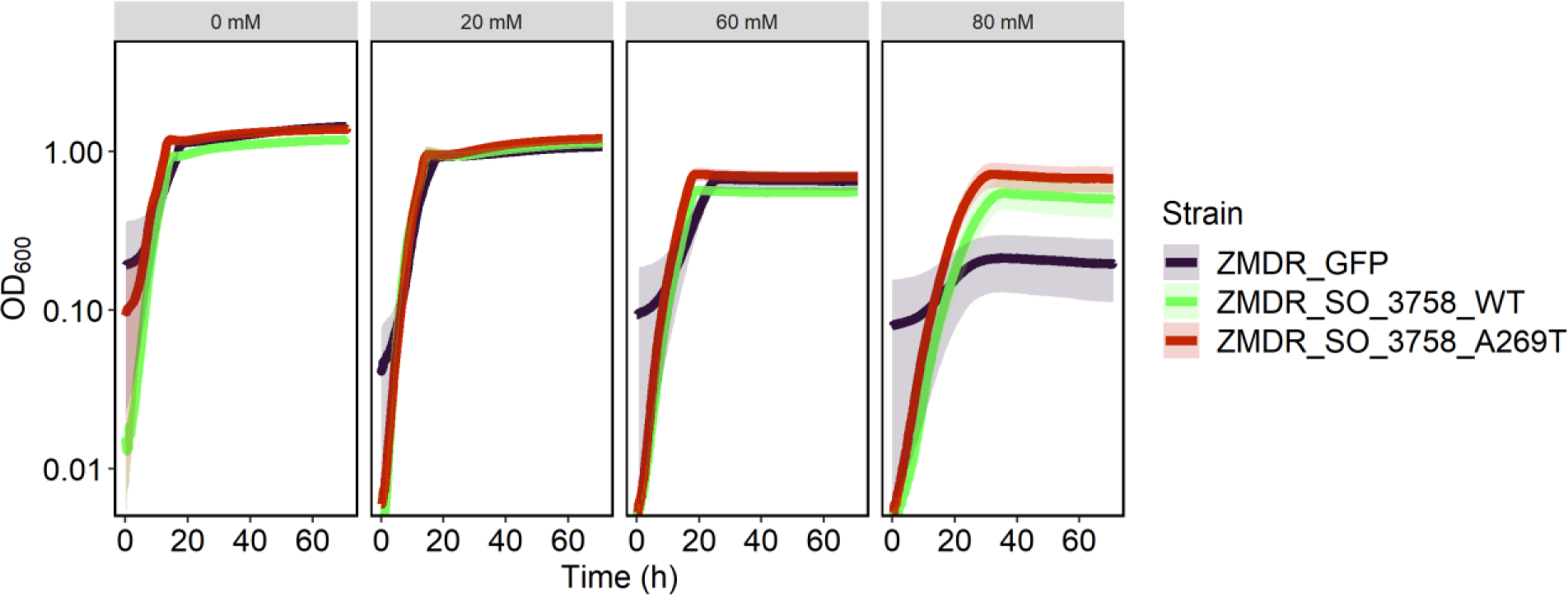
Growth curves of *Z. mobilis* ZMDR harboring an IPTG-inducible expression vector containing GFP, SO_3758_WT or SO_3758_A269T in 200 μl ZRMG medium with varying formate concentrations.

The effects of mutations in DUF2721-containing protein SO_1320 are more difficult to ascertain. While the function of both DUF2721 and SO_1320 are unknown, some clues exist to the protein’s potential role. Both YadS and SO_1328 expressed from the same operon as SO_1320 are part of membrane protein families. YadS is annotated as a trimeric intracellular cation (TRIC) channel and SO_1318 is annotated as a nuclear transport factor 2-like (NTF2) protein (Heidelberg et al., 2002). The proximity of SO_1320 to these membrane-bound transporter proteins could be an indicator that it shares a similar function.

In an attempt to elucidate the potential function of SO_1320, the amino acid sequence was analyzed using DeepGO, a deep learning tool which predicts localization and function of a protein based on its sequence (Kulmanov et al., 2018). DeepGO predicted that SO_1320 is localized to the cell membrane but was unable to predict its function, likely due to a lack of experimentally characterized homologs (**Table S4**). Further analysis of the SO_1320 amino acid sequence by DeepTMHMM (Hallgren, Tsirigos, Pedersen, Armenteros, et al., 2022), a tool for predicting transmembrane proteins using deep neural networks, reiterates the protein’s probable localization to the cell membrane, finding three transmembrane helices (**Figure S5**). These results suggest that

SO_1320 is associated with the cellular membrane, although it is still unclear what function it serves. Because the formate concentrations are constant throughout culturing of MGC003, MGC004, and MGC007 (formate tolerant strains with mutations in SO_1320), the evolved method of formate tolerance must not involve the oxidation or assimilation of formate. Potential formate tolerance mechanisms of SO_1320 mutants could include enhanced active formate efflux, decreased active formate influx, or membrane stabilization preventing passive formate diffusion.

To determine if SO_1320 could bind formate, the unmutated and mutated versions of SO_1320 were modeled using AlphaFold (Jumper et al., 2021) and formate binding was predicted with AutoDock Vina (Trott & Olson, 2010). SO_1320 did not appear to have convincing formate binding activity, as the best predicted binding affinity was -2.1 kcal mol^-1^ at a site on the protein surface (**Figure S6**). Modeling of formate binding to SO_1320_V106I, SO_1320_V106F, and SO_1320_S52I calculated predicted binding affinities to formate of -2.0 kcal mol^-1^, -2.1 kcal mol^-1^, and -2.1 kcal mol^-1^, respectively. The low predicted binding affinity, the location of the predicted binding, and the lack of improvement to the predicted binding affinity in the mutants suggest that neither SO_1320 nor its mutants are acting as a formate transporter; however, it does not eliminate the possibility that it could be a formate channel or that it could form a multimeric complex with a formate transporter whose substrate binding does not involve SO_1320 directly.

## Discussion

This work describes the evolution of seven formate-tolerant *S. oneidensis* strains for use in systems subjected to high formate concentrations. Two notable mutations arose during evolution of *S. oneidensis* for greater formate tolerance. The first is a mutation in alanine 269 to a threonine in the sodium-dependent bicarbonate transporter SO_3758. Every strain exhibiting this mutation in SO_3758 (MGC002, MGC005, and MGC006) shared the same A269T substitution and was able to withstand greater formate concentrations than the parent strains. The second gene with mutations attributed to greater formate tolerance was SO_1320 encoding a protein of unknown function. Mutations occurred in residue 106, changing the native valine to either isoleucine (MGC007) or phenylalanine (MGC003 and MGC004), and in residue 52 (MGC008), changing the native serine to isoleucine. The evolved strains gained the capacity to grow robustly on 100 mM formate in minimal lactate medium when the parent strains they are derived from were only capable of growth on 10 mM formate.

Strains harboring a mutation in V106 of SO_1320 had a shorter lag phase than strains harboring the A269T mutation in SO_3758, although all strains grew to a similar final OD_600_ (**Figure 3**). Therefore, the onset of formate tolerance benefits from the SO_1320 mutations occurred earlier than those from the SO_3758_A269T mutation. The extra delay in growth of strains with the A269T mutation in SO_3758 may be related to the mechanism by which it provides formate tolerance. Modeling of formate binding in the WT and mutant versions of SO_3758 revealed that the conversion of alanine 269 to threonine increased the binding affinity of formate to a site deeper in the binding pocket. This site is predicted to bind bicarbonate in the native protein with a lower predicted binding affinity, indicating that the mutation did not create a new binding site, but rather strengthened an existing one. The predicted binding affinity of both bicarbonate and formate in the deeper binding site of SO_3758_A269T is stronger than the binding of both substrates to both the native and deeper binding site in the unmutated protein. The mutation allows formate binding to a similar efficiency of the native bicarbonate binding, suggesting that the mutation has made formate a suitable substrate for SO_3758_A269T. The ability to better bind formate suggests that the strains harboring this mutation had improved formate efflux capabilities over the WT. Without the benefit of being able to transport formate, the parent strains with WT SO_3758 were less efficient at preventing the toxic accumulation of formate in the cells.

Further evidence for the formate binding capabilities of both the WT and mutant versions of SO_3758 is observed during their expression in *Z. mobilis*. Interestingly, SO_3758_WT confers similar formate tolerance as SO_3758_A269T in this organism, an observation that differs from their performance in *S. oneidensis*. *Z. mobilis* does not have a version of the Sbt-like sodium-dependent bicarbonate transporter (Seo et al., 2005). This may suggest that the introduction of even the WT SO_3758, which binds formate weaker than SO_3758_A269T, can provide an effective mechanism for formate efflux that is otherwise absent. The ability to transfer improvement of formate tolerance to other species through expression of a single protein may be useful in engineering a variety of bacterial biosynthetic hosts that are limited by formate toxicity. Additionally, further protein engineering could improve on the single A269T mutation and create an even stronger formate transporter that could further improve formate tolerance.

The role of mutations in DUF2721-containing protein SO_1320 are less clear as neither this domain nor this protein have any known characterized homologues. SO_1320 is in an operon with genes encoding a membrane protein channel or membrane transport protein (*yadS* and SO_1318), and sequence analysis tools predict three transmembrane helices. These clues suggest that SO_1320 is also a membrane protein. Modeling of formate binding to WT and mutant versions of SO_1320 indicate that the protein does not bind formate as a transporter might. While it is still unclear how SO_1320 mutants are providing cells with greater formate tolerance, our analysis does not rule out the possibilities that SO_1320 is involved in membrane stabilization or acts as a formate channel. Additionally, SO_1320 could be one subunit in a multimeric formate transporter which binds formate in a different subunit.

Expression of these mutant genes in the *S. oneidensis* WT background on minimal lactate medium increased formate tolerance two-fold. The ability to reconstruct the formate tolerant phenotype through expression of these mutated proteins further confirms their role in the evolved strains. The simple point mutations which lead to these growth improvements provide two easily accessible methods for transferring formate tolerance to *S. oneidensis* and other organisms. This could be a valuable tool in engineering biocatalysts that would benefit from reduced intracellular formate or bicarbonate levels, such as in organisms in which these compounds block metabolic activity prematurely due to toxicity.

## Corresponding Author

Michaela A TerAvest – ORCID: 0000-0002-5435-3587;

## Authors

Megan C Gruenberg Cross – ORCID: 0000-0002-9158-9900; Elhussiny Aboulnaga – ORCID: 0000-0002-8664-9497;

## Data Availability

Raw sequencing reads are available upon request.

## Acknowledgements

We thank Dr. Jeffrey Gralnick for *S. oneidensis* strains JG2955 and JG2957. We also thank Drs. Nanye Long and Nicholas Panchy for their assistance with Breseq and AlphaFold. We acknowledge support from the Michigan State University Bioinformatics Core.

